# Carrier-set topology reveals composite MHC tagging at the MICA rs2596542 locus: insights from a flip-flop association example

**DOI:** 10.64898/2026.05.15.725353

**Authors:** Yuki Ichikawa

## Abstract

Cross-population reversal of signed linkage disequilibrium (LD), or the “flip-flop” phenomenon, can arise when a tag SNP captures different extended haplotype backgrounds across populations. The MICA hepatocellular carcinoma susceptibility variant rs2596542 exemplifies this problem in the MHC, where signed LD reverses between Japanese and European populations but the relevant regulatory backgrounds are obscured by haplotypic complexity. We analyzed 7,303 biallelic SNVs surrounding rs2596542 across 26 populations using carrier-set topology classification followed by non-negative matrix factorization of carrier haplotypes. This identified two regulatory axes. Axis I, represented by components c4/c6, was population-stable and MICA-regulatory, with coherent MICA cis-eQTL enrichment and depletion for signed-LD reversal. Axis II, represented by component c5, was enriched for signed-LD reversal and showed an HLA-B↑/HLA-C↓ expression signature with no MICA overlap across six GTEx tissues. The previously proposed cross-population tag rs2244546 mapped to a population-stable component rather than Axis II. Together, these findings indicate that rs2596542-T is not a single MICA-regulatory proxy but a composite MHC tag that captures separable MICA- and HLA-B/HLA-C-regulatory haplotypic backgrounds. Carrier-set topology combined with NMF provides a practical approach for resolving such composite tag signals and for interpreting cross-population signed-LD reversal in complex MHC loci.

## Introduction

The major histocompatibility complex (MHC) harbors extensive haplotypic diversity shaped by balancing selection, recombination suppression, and population-specific evolutionary pressures. This complexity creates a central challenge for genetic association studies: tag SNPs selected in one population may capture different functional backgrounds in another, because the same marker allele can reside on structurally distinct extended haplotypes whose proportions vary across populations.

A specific manifestation of this problem is the “flip-flop” phenomenon, described systematically(Lin et al. 2007), in which a GWAS-identified risk allele shows opposite effect directions across populations as a consequence of cross-population variation in LD between observed marker alleles and unobserved causal variants. Subsequent theoretical work has shown that flip-flop can also arise through population shifts in haplotype-frequency configuration, even when parts of the underlying LD structure are conserved(Zaykin and Shibata 2008). Under simple penetrance models, such shifts at proxy SNPs can be formalized as reversals in the sign of LD between marker and causal alleles(Clarke and Cardon 2010). Together, these frameworks motivate a composite-tagging perspective, in which a single tag SNP captures multiple functionally distinct extended haplotypic backgrounds whose mixture proportions vary across populations. Despite its theoretical prominence, this mechanism has rarely been resolved empirically at specific disease-associated loci.

The MICA variant rs2596542 exemplifies this problem. It was first identified as a hepatocellular carcinoma (HCC) susceptibility locus in a Japanese GWAS(Kumar et al. 2011), and subsequent work linked this association to a promoter variant influencing MICA expression(Kumar et al. 2012; Lo et al. 2013). However, rs2596542 has shown inconsistent effect directions across populations: opposite associations have been reported in European cohorts(Augello et al. 2018), and meta-analyses support population heterogeneity(Kuang et al. 2019). Lange et al. further showed that signed Pearson correlations between rs2596542 and nearby MHC markers reversed between Japanese and European populations, and proposed rs2244546 near HCP5 as a more stable cross-population tag(Lange et al. 2013). These observations suggest that rs2596542 is not merely a local MICA-expression tag, but a composite marker whose population-specific signal may depend on the mixture of extended MHC haplotypes it captures. Yet existing analyses have largely focused on individual MICA-flanking variants or pairwise LD with a limited number of partners. The broader LD architecture surrounding rs2596542 — spanning HLA-B, HLA-C, MICB, and adjacent regulatory regions — has not been jointly evaluated as an integrated haplotype space.

To address this gap, we applied a two-step framework. First, we classified partner SNVs by carrier-set topology (DISJOINT, PARTIAL, and STRICT inclusion classes) to determine whether signed-LD reversal reflected discrete carrier-set boundary changes or continuous mixture shifts within a shared carrier space. We found that reversal was concentrated within PARTIAL-overlap topology rather than DISJOINT or STRICT boundary classes, indicating that the relevant variation was not a simple gain or loss of carrier-set overlap but a shift in the mixture of haplotypic backgrounds carried by rs2596542-T. We therefore applied non-negative matrix factorization (NMF) to rs2596542-T carrier haplotypes across 26 populations to decompose this shared carrier space into latent haplotypic signatures. This decomposition resolved two functionally distinct regulatory axes: a population-conserved MICA cis-regulatory axis and a population-variable HLA-B/HLA-C axis enriched for cross-population signed-LD reversal. We further evaluated these axes using cis-eQTL data across multiple tissues.

## Methods

### Data sources

rs2596542 alleles are given on the forward strand; rs2596542-T corresponds to the A allele reported by Kumar et al(Kumar et al. 2011). Variant coordinates are reported in GRCh37/hg19 unless otherwise stated, with allele identities harmonized to the GRCh38/dbSNP forward strand. Phased haplotypes from 1000 Genomes Phase 3 (The 1000 Genomes Project Consortium 2015) for 26 populations. All biallelic SNVs within ±200 kb of rs2596542 (chr6, GRCh37) with MAF ≥ 0.01 in at least one super-population were retained (7,303 variants). The anchor variant rs2596542 (chr6:31,366,595, GRCh37) is encoded in the 1000 Genomes Phase 3 VCF as C reference and T alternate on the forward strand; haplotype value 1 therefore corresponds to the rs2596542-T-carrying haplotype. Carrier haplotypes: rs2596542-T allele carriers on the forward strand. Within rs2596542-T carrier haplotypes, SNVs were retained only if their carrier-subset ALT frequency was between 0.05 and 0.95, excluding both rare and near-fixed alleles that contribute little to haplotype decomposition. The anchor SNV rs2596542 itself was removed from the NMF input matrix after carrier selection because all retained haplotypes carry the ALT allele at this site, leaving it with zero variance in the analytic matrix. GTEx v8 cis-eQTL summary statistics(GTEx Consortium 2020) were obtained for six tissues: Whole Blood, Liver, Lung, Sun-Exposed Skin, EBV-transformed lymphocytes, and Transverse Colon.

### Pairwise LD and signed reversal

For each anchor–partner pair, we computed signed Pearson r, the classical LD coefficient D, and a derived asymmetry component C = D·(1/p_B − 1/p_A) per population. C is not introduced here as a new LD statistic; rather, it is an algebraic descriptor of the LD-coupled term in the conditional-probability difference P(A|B) − P(B|A). Although conditional asymmetric LD has been formalized previously for biallelic markers(Thomson and Single 2014), the present analysis does not use ALD itself; instead, C is used only to decompose the directionality of P(A|B) − P(B|A).

Specifically, P(A|B) − P(B|A) decomposes exactly into a marginal allele-frequency term, p_A − p_B, and an LD-coupled term, C = D·(1/p_B − 1/p_A). Because sign(C) = sign(D)·sign(p_A − p_B), C is sensitive not only to reversals of LD sign, as captured by signed r, but also to reversals of marginal allele-frequency rank under conserved D. Comparing signed-r reversal and C reversal therefore separates LD-sign changes from frequency-rank changes, providing a way to relate the proxy-LD mechanism emphasized by Clarke and Cardon(2010) to constant-LD flip-flop configurations such as those described by Zaykin and Shibata(2008).

Signed-r reversal was defined as a sign change in signed Pearson r between any EAS and EUR population pair. C reversal was defined analogously as a sign change in C. Carrier-set topology was classified as DISJOINT (n_11_ = 0), STRICT inclusion (n_10_ = 0 or n_01_ = 0, with n_11_ > 0), or PARTIAL overlap (all four haplotype-count cells > 0). C was included to test whether signed-r reversal reflected continuous directional LD change rather than discrete carrier-set topology transitions.

### UMAP and Leiden clustering

675 × 7,303 binary haplotype matrix embedded with UMAP (n_neighbors = 15, min_dist = 0.1, metric = hamming). Leiden community detection on the k-nearest-neighbor graph (resolution = 1.0).

### Non-negative matrix factorization

NMF was used as an interpretable signature-discovery method rather than as a formal population-ancestry model. Similar matrix-factorization approaches have been widely used for signature discovery in genomics, including gene-expression decomposition and mutational signature analysis(Brunet et al. 2004; Alexandrov et al. 2013). Although ancestry-decomposition methods such as ADMIXTURE(Alexander et al. 2009) also represent samples as non-negative mixtures, they assume an allele-frequency likelihood in which components correspond to ancestral populations. This was not the objective here. We focused on haplotypes carrying the same rs2596542-T anchor allele and sought to identify co-occurring SNV patterns within that carrier set that could be directly annotated by LD reversal, genomic location, and cis-eQTL effects. NMF provides both haplotype loadings (W matrix) and component-defining SNV signatures (H matrix), making it suitable for this purpose. Accordingly, NMF components are interpreted as latent haplotypic SNV signatures, not as ancestral populations or phylogenetic lineages.

NMF was performed using scikit-learn (sklearn.decomposition.NMF) with Frobenius norm loss, NNDSVD initialization, a maximum of 500 iterations, and tolerance 10^−4^. NMF was run across ranks k = 4–12. Consensus-clustering cophenetic correlation across k = 2–12 with 20 random seeds per rank placed k = 8 (cophenetic = 0.963) within 0.05 of the maximum while resolving the MICA-stable, HLA-B-stable, and HLA-B/HLA-C-variable signatures separately; k = 8 was therefore selected as the primary rank.

Fifty-seed reproducibility at k = 8 confirmed high stability: mean Hungarian-matched H-row correlations were 0.98, 0.96, and 0.99 for c4, c5, and c6, respectively, with MICA-enriched and HLA-B-enriched component labels recovered in 94% and 100% of seeds. Component sparsity was quantified by the Gini coefficient of each component’s H and W vectors. Top-signature SNVs were defined as those with H-matrix loadings in the top 5% of each component (≥95th percentile), with sensitivity analysis across top 1–15% thresholds.

### GTEx eQTL direction analysis

For each component and target gene, sign coherence and NES distribution (Wilcoxon rank-sum test) were computed across six tissues.

### Analysis structure

The c5 axis was defined exclusively in 1000 Genomes Phase 3 normal haplotype space using LD reversal, NMF decomposition, and GTEx eQTL annotation. No tumor-expression or disease-outcome data were used to define or refine the component structure.

## Results

### Cross-population signed-LD reversal at rs2596542

We computed pairwise signed Pearson r (and the underlying signed D) between rs2596542 and 7,303 biallelic SNVs within ±200 kb across 26 populations from 1000 Genomes Phase 3. In parallel, we additionally tracked C = D·(1/p_B − 1/p_A), a covariance-derived descriptor used here to separate LD-sign reversal from allele-frequency-rank reversal (see Methods). Of these partner SNVs, 33% showed signed-r reversal between East Asian (EAS) and European (EUR) population pairs, increasing to 59% when African (AFR) populations were included. C reversal showed high overlap with signed-r reversal (Jaccard = 0.755). This overlap indicates that cross-population reversal at this locus is dominated by LD sign change rather than allele-frequency-rank reversal (see Methods for the signed-r vs. C decomposition; cf. Clarke and Cardon(2010); Zaykin and Shibata(2008)).

Carrier-set topology class changes (transitions between DISJOINT, PARTIAL, and STRICT inclusion classes) occurred at 13–14% of partner SNVs with minimal overlap with continuous r/C reversal (Jaccard = 0.13–0.14), establishing two distinct regimes of cross-population LD change: continuous signed reversal within the PARTIAL class and discrete topology transitions at haplotype boundaries.

**Figure 1.**
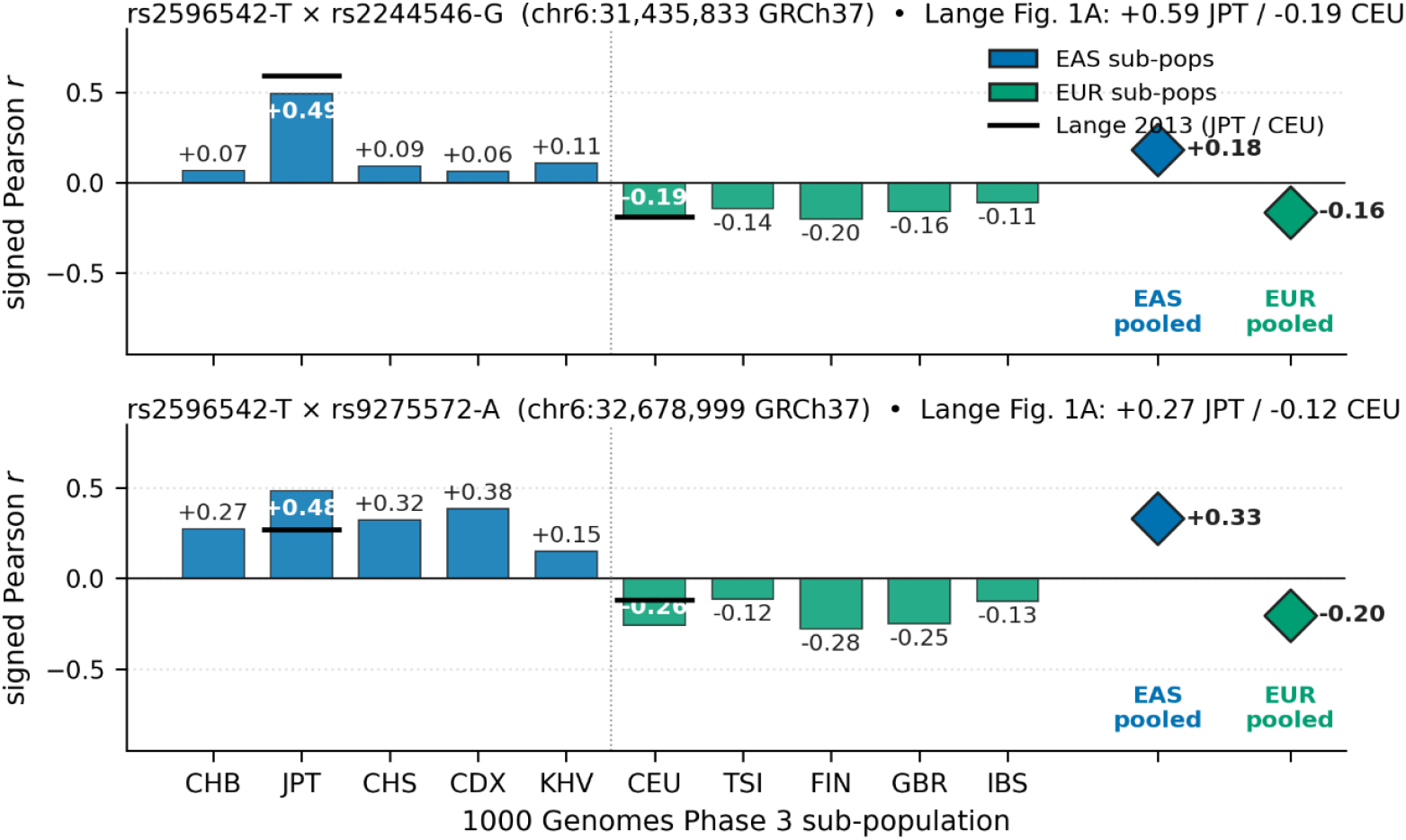
Population-conditional sign of LD between rs2596542 (anchor) and two HCV-HCC tag candidates — Lange 2013 Fig. 1A replication in 1000G phase 3. Cross-population signed-LD reversal at MICA rs2596542. Signed Pearson r for rs2596542 paired with rs2244546 (top) and rs9275572 (bottom) across EAS and EUR populations, replicating Lange et al. (2013). Black horizontal bars at JPT and CEU positions show Lange’s(Lange et al. 2013) published values. Diamonds at right show pooled EAS and EUR signed Pearson r computed by combining haplotypes across sub-populations within each super-population.

### Latent haplotype branch decomposition

UMAP dimensionality reduction and Leiden community detection applied to 675 rs2596542-T carrier haplotypes from 10 EAS/EUR populations identified 19 clusters with strong population stratification. Several clusters were exclusive to East Asian haplotypes (L0, n = 60; L9, n = 37), others were European-specific (L7, Finnish-enriched, n = 41; L13, Southern European, n = 23), and one was private to Japanese samples (L18, n = 5). Extension to 26 populations revealed five additional African-private clusters, including Luhya-dominant and Mende-dominant haplotypic backgrounds.

NMF was performed at k = 8, selected by consensus cophenetic stability (0.963 across 20-seed runs over k = 2–12) as the smallest rank that separately resolved the MICA-stable, HLA-B-stable, and HLA-B/HLA-C-variable signatures (see Methods; Supplemental Fig. S2). At this rank, NMF decomposed each carrier haplotype into a weighted combination of eight latent components. Gini coefficient-based sparsity analysis classified components along a spectrum from branch-specific (high sparsity: c4, Gini = 0.90) to backbone (lower sparsity: c7, Gini = 0.78).

**Figure 2.**
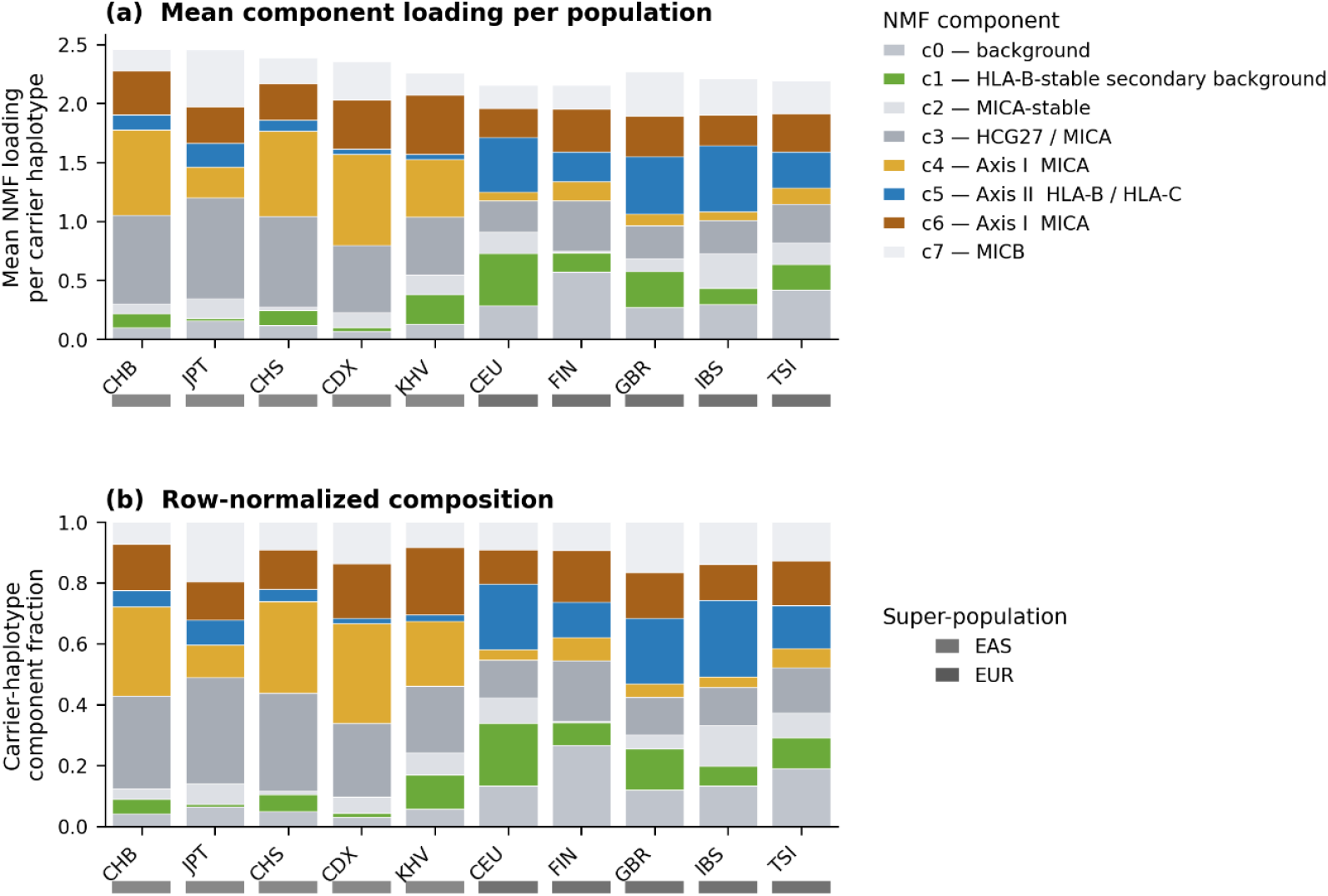
k = 8 NMF component composition among rs2596542-T carrier haplotypes in EAS and EUR populations. Stacked bars show mean NMF component loading and row-normalized NMF component composition within rs2596542-T carrier haplotypes for EAS and EUR populations. Components c4 and c6 correspond to the Axis I MICA cis-regulatory background, whereas c5 corresponds to the Axis II HLA-B/HLA-C regulatory background enriched for signed-LD reversal. Super-population membership is indicated below each population. This k = 8 decomposition shows that rs2596542-T carriers comprise a mixture of latent haplotypic backgrounds whose proportions vary across populations.

### Two functionally distinct regulatory axes

Axis I (MICA cis-regulatory axis). Components c4 and c6 were enriched for MICA cis-eQTL SNVs from GTEx Whole Blood (c6: fold = 5.58×; c4: fold = 3.29×; both P < 10^−15^, the display cap) with gene-body enrichment within MICA and MICA-AS1 (fold 9.2–9.3×). These components were depleted for signed-r reversal (c6: fold = 0.13) and C reversal (c6: fold = 0.73), but enriched for topology class changes (c6: fold = 2.15×). This axis captures population-conserved MICA regulatory variation.

Axis II (HLA-B/HLA-C regulatory axis). Component c5 showed the opposite profile: enrichment for signed-r reversal (fold = 1.48×) and C reversal (fold = 1.27×), depletion for topology class changes (fold = 0.00), and a region-wide diffuse SNV signature. c5 was enriched for HLA-B eQTLs (fold = 2.14×) and HLA-C eQTLs (fold = 2.49×; both P < 10^−15^, the display cap) but showed zero MICA eQTL overlap at all thresholds tested (fold = 0.00, top 1–15%).

A third component (c1, HLA-B-stable) was enriched for HLA-B eQTLs (fold = 2.98×; P < 10^−15^, the display cap) but showed no LD reversal enrichment (C-flip fold = 1.02), demonstrating that HLA-B regulation has both population-stable and population-variable components, with only the latter contributing to cross-population signed-LD reversal. Component c1 showed lower seed stability than the primary MICA-stable (c4) and HLA-B/HLA-C-variable (c5) components (Supplemental Fig. S3) and is interpreted here as a secondary HLA-B-associated background that supports the contrast with c5 rather than as a fully reproducible primary axis.

### Expression direction coherence across tissues

Axis I MICA-down direction is conserved across tissues. All c4 and c6 signature SNVs with MICA cis-eQTLs showed negative effect sizes across six GTEx tissues (Whole Blood, Liver, EBV-transformed lymphocytes, Lung, Skin Sun-Exposed, Colon Transverse), with sign coherence = 1.00 in every tissue–component combination (c4: 1,010 SNV-tissue observations, n_neg/n_pos = 1,010/0; c6: 1,721 observations, 1,721/0; combined 2,731/0). Mean NES magnitude was tissue-dependent, strongest in Colon (c4: −0.72; c6: −0.67) and Skin (−0.61; −0.55) and weakest in Whole Blood (−0.37; −0.35), consistent with tissue-specific MICA regulatory amplitude.

All c5 signature SNVs with HLA-B eQTLs showed positive effects in Whole Blood (229/0, sign coherence = 1.00), all with HLA-C eQTLs showed negative effects (0/296, coherence = 1.00), yielding a consistent HLA-B↑/HLA-C↓ pattern (Wilcoxon P: HLA-B = 10^−39^, HLA-C = 10^−50^). This directional pattern replicated across six GTEx tissues (HLA-B detected in five; EBV-transformed lymphocytes lacked HLA-B eQTL overlap). In liver, c5 showed HLA-B↑ (52/0), HLA-C↓ (0/273), and HCG27↑ (228/0), with effect sizes equal to or stronger than blood (HLA-C NES: −0.62 vs −0.25). In colon, the same pattern held (HLA-B↑ 134/0, HLA-C↓ 0/296), although with HLA-C magnitude substantially weaker than in liver (NES = −0.24 vs −0.62), indicating that the c5 HLA-C effect is most pronounced in liver. MICB modulation was tissue-dependent (positive in blood, absent in liver, negative in skin), indicating that the core c5 signature is HLA-B↑/HLA-C↓/HCG27↑ with tissue-specific MICB co-regulation.

Two distinct HLA-B-upregulating haplotypic backgrounds coexist within the rs2596542-T carrier set: c1 (HLA-B↑/MICB↓/HLA-C mixed, population-stable) and c5 (HLA-B↑/HLA-C↓/MICB↑, population-variable), representing different extended MHC haplotypic backgrounds that share HLA-B upregulation but differ in HLA-C and MICB co-regulatory direction.

**Figure 3.**
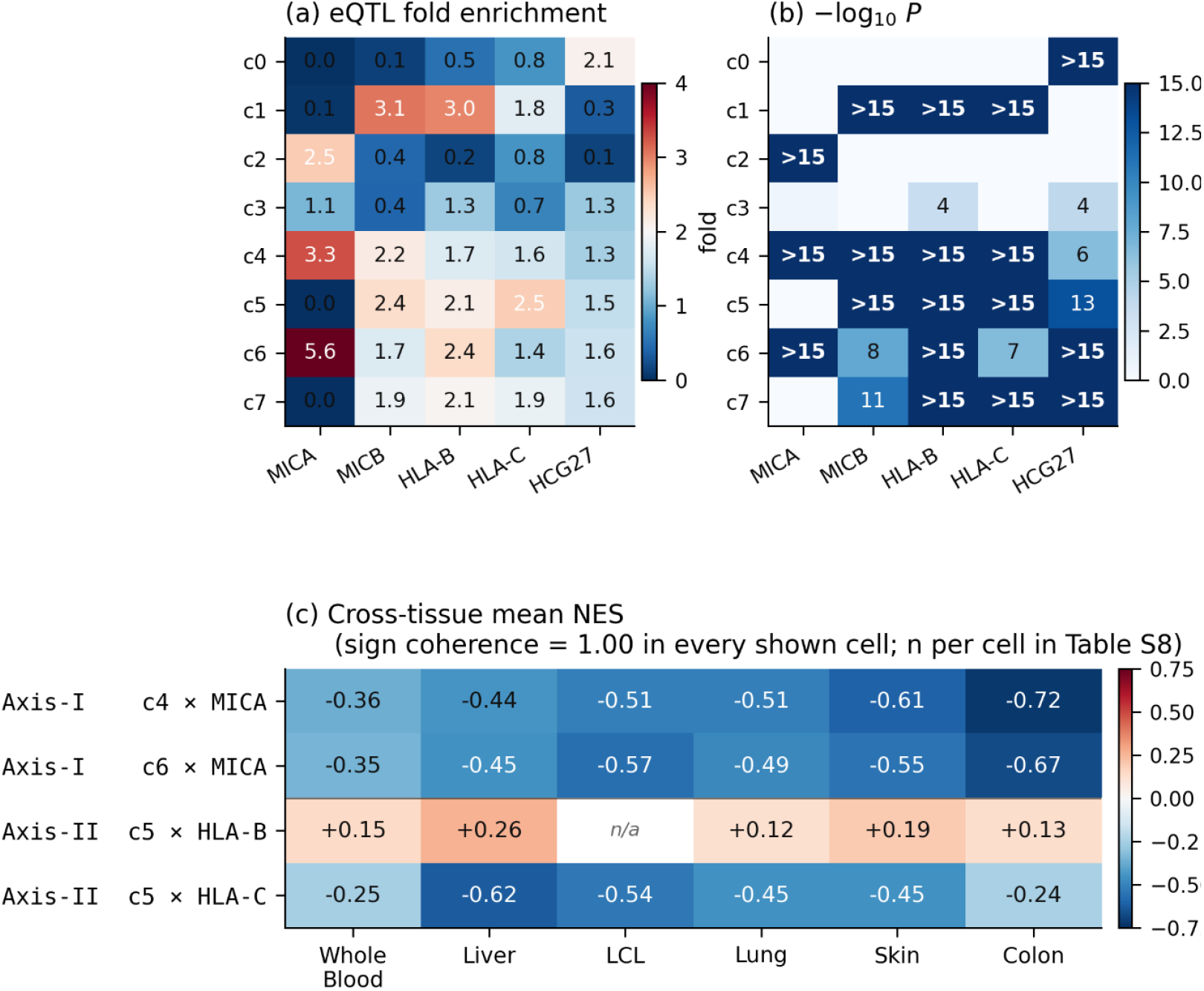
NMF components separate MICA↓ and HLA-B↑/HLA-C↓ regulatory axes. NMF component × target-gene eQTL correspondence. (a) Fold enrichment of component top-5% SNVs for cis-eQTLs of each gene. c4/c6 define Axis I (MICA-enriched, MICA↓); c5 defines Axis II (HLA-B/HLA-C-enriched, HLA-B↑/HLA-C↓, with MICA = 0). (b) Hypergeometric significance (−log_10_ P, capped at 15). HCP5 column shown as n/a in both panels due to insufficient overlap with the top-5% SNV set for enrichment calculation. (c) Cross-tissue mean NES (GTEx v8) for the principal axis–gene combinations. Sign coherence = 1.00 in every shown cell (c5 × HLA-B × LCL had no eQTL overlap and is shown as n/a). Axis I (c4/c6 × MICA) shows monotonically increasing magnitude from Whole Blood (mean NES ≈ −0.35) to Colon (≈ −0.70); Axis II HLA-C downregulation is strongest in Liver (mean NES = −0.62). Combined: Axis I, 2,731 SNV-tissue observations all NES < 0; Axis II, 921 HLA-B↑ + 1,698 HLA-C↓ = 2,619 observations all direction-consistent. Full per-tissue counts in Supplemental Table S8.

### Tag SNPs map to stable axes

Among previously reported tag SNPs at this locus, rs2395029 (HCP5/HLA-B*57:01 proxy) and rs2244546 were directly present in the NMF SNV set, while rs2596542 had been removed from the NMF input matrix after carrier selection and was represented by its nearest in-window NMF SNV. Their H-matrix loadings placed rs2395029 in c1 (HLA-B-stable), whereas rs2596542 and rs2244546 mapped to c3 (mixed). None of these tag SNPs or proxies fell within the top 5% of any component, indicating that the HLA-B/HLA-C-variable axis (c5) represents a co-occurring SNV signature not captured by existing tags.

### Robustness analyses

The two-axis structure was robust to NMF rank (k = 4–12: both axes present from k = 5, with consensus cophenetic correlation 0.963 at k = 8), signature threshold (top 1–15%: c5 MICA fold = 0.00 at all thresholds), random initialization (50-seed reproducibility: mean matched correlations 0.96–0.99 for c4/c5/c6), and MAF/distance-matched permutation testing (10,000 permutations: c5 r-flip z = +10.8, C-flip z = +11.3, HLA-B z = +10.9, HLA-C z = +13.3). A parallel probabilistic decomposition using Latent Dirichlet Allocation recovered the major component structure (matched correlations 0.87–0.99 for c4/c5/c6) but attenuated the localized HLA-B eQTL enrichment, consistent with smoothing of sparse regulatory signatures by mixed-membership models (Supplemental Figs. S1–S3).

**Figure 4.**
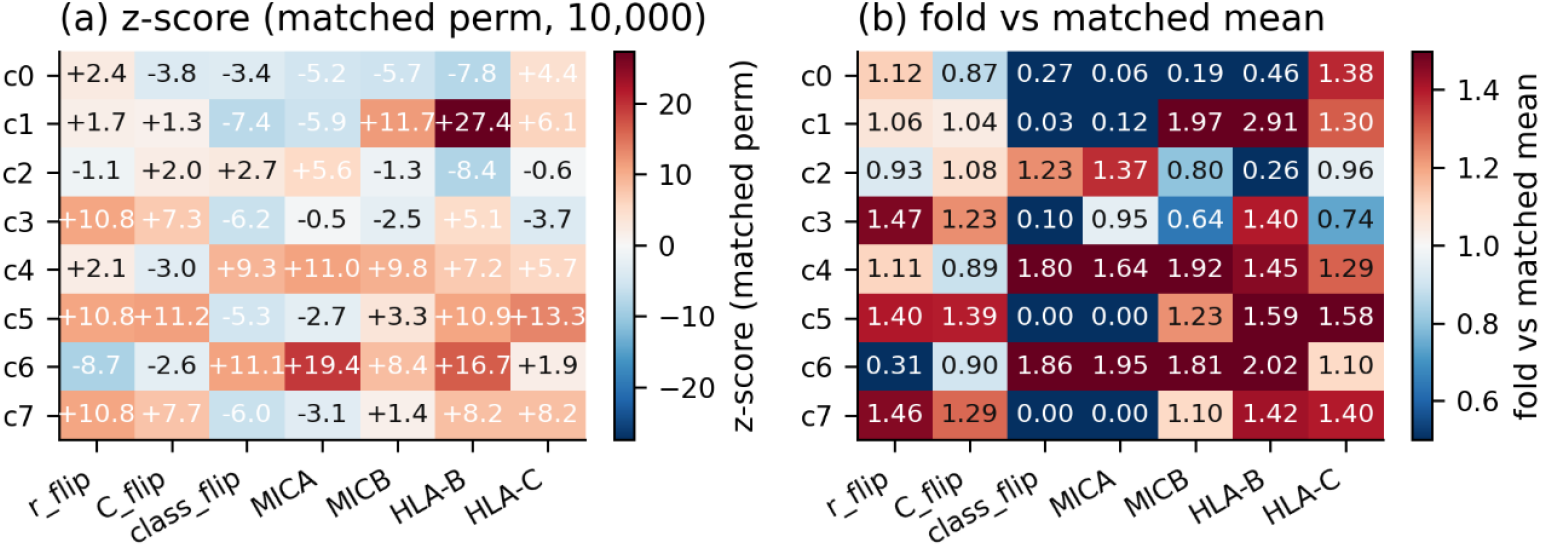
MAF×distance matched permutation (k = 8, top 5 %, 10,000 perms). MAF × distance-matched permutation robustness (10,000 permutations). (a) z-scores: c5 is strongly enriched for r-flip, C-flip, HLA-B, HLA-C and depleted for MICA; c4/c6 show the complementary pattern. (b) Fold enrichment vs matched null mean.

## Discussion

### Functional decomposition of the rs2596542-T carrier space

We show that rs2596542-T carrier haplotypes in the MHC resolve into two functionally distinct regulatory axes with different population dynamics and gene-regulatory targets. Axis I, represented by components c4 and c6, corresponds to a population-conserved MICA cis-regulatory axis. This axis was enriched for MICA cis-eQTLs, with the MICA-down direction reproduced across all six GTEx tissues tested. Axis II, represented by component c5, corresponds to a population-variable HLA-B/HLA-C regulatory axis. This axis carried a coherent HLA-B↑/HLA-C↓ expression signature and was enriched for cross-population signed-LD reversal.

These two axes also showed distinct tissue-magnitude profiles. Axis I MICA downregulation was stronger in colon and skin than in whole blood, whereas Axis II HLA-C downregulation was strongest in liver. This liver-pronounced HLA-C regulatory signature provides a plausible context for considering the HLA-B/HLA-C axis in HCC-associated MHC tagging. More broadly, these findings indicate that rs2596542-T is not a single functional tag for MICA, but a composite carrier state containing separable MICA-regulatory and HLA-B/HLA-C-regulatory haplotypic backgrounds.

### Composite tagging as a mechanism for rs2596542 flip-flop

The cross-population signed-LD reversal observed at rs2596542 is consistent with a shift in the mixture proportions of functionally distinct MHC regulatory haplotypes, rather than reversal of a single causal effect. Within the rs2596542-T carrier set, at least two HLA-B-regulating backgrounds coexist: one population-stable and one population-variable. When the frequency of the c5-tagged haplotypic background shifts across populations, the net signed LD between rs2596542 and downstream markers can change direction. This provides a candidate haplotypic mechanism for the reported flip-flop associations at this locus.

This interpretation extends the observation of Lange et al., who proposed rs2244546 near HCP5 as a more stable cross-population tag(Lange et al. 2013). Our component mapping supports the stability of this tag but clarifies its scope: rs2244546 mapped to a population-stable component rather than the population-variable HLA-B/HLA-C axis. Thus, rs2244546 may capture part of the conserved MHC background around rs2596542, but it does not resolve the specific c5 axis whose population-dependent frequency shift is expected to contribute to signed-LD reversal.

### Potential relevance to HCC susceptibility and flip-flop association

rs2596542 was originally identified as an HCC susceptibility locus and has largely been interpreted through MICA-centered biology. The present component-level analysis refines this interpretation by showing that rs2596542-T marks not only a population-conserved MICA-regulatory axis but also a population-variable HLA-B/HLA-C-regulatory axis. Because HLA-C is a key ligand in NK-cell receptor biology and HLA class I dysregulation has been implicated in HCC tumor biology and prognosis(Wang et al. 2019; Hazini et al. 2021; Pei et al. 2022), this HLA-B/HLA-C axis provides a biologically plausible haplotypic background that may contribute to the population-dependent interpretation of rs2596542. Thus, the reported flip-flop behavior at this locus may reflect changes in the mixture of MICA-centered and HLA-C-linked regulatory backgrounds rather than reversal of a single MICA-mediated effect. Direct mediation of HCC susceptibility by the HLA-B/HLA-C axis remains to be tested in ancestry-diverse genetic and clinical cohorts.

### Composite tagging as a haplotypic heterogeneity class

The mechanism documented here can be viewed as a haplotypic form of composite-tagging heterogeneity. It differs from classical allelic heterogeneity, in which different alleles at the same locus drive association in different populations, and from locus heterogeneity, in which different loci are responsible. Here, the same marker allele tags structurally distinct extended haplotypic backgrounds, and cross-population changes in their mixture proportions alter the observed LD and regulatory signal.

This framework helps reconcile two views of flip-flop association. Proxy-LD models emphasize reversal in the sign of LD between marker and causal alleles, whereas haplotype-frequency models show that effect-direction reversal can arise through population shifts in haplotype configuration even when parts of the LD structure are conserved. The rs2596542-T carrier space provides an empirical example in which these mechanisms can be connected: component-level decomposition reveals how a shared marker allele can carry multiple regulatory backgrounds whose mixture proportions differ across populations.

### Implications for MHC association studies

These results raise the possibility that reported rs2596542 associations partly reflect the HLA-B/HLA-C regulatory background tagged by this SNP, rather than MICA coding or expression effects alone. This does not exclude a role for MICA biology. Instead, it suggests that the rs2596542 association should be interpreted as a composite MHC haplotypic signal rather than as a simple single-gene marker.

More generally, the analytical framework used here—carrier-set topology classification followed by NMF decomposition of carrier haplotypes and functional annotation using LD reversal and cis-eQTL enrichment—may be useful for resolving other population-inconsistent association signals in the MHC. Such cases are often difficult to interpret using pairwise LD or single-SNP proxy substitution alone, because the same marker allele may tag different extended haplotypic backgrounds in different populations. Component-aware carrier-haplotype analysis provides a way to decompose these composite signals into functionally interpretable subspaces.

### Limitations and future directions

Several limitations should be considered. First, NMF is a model-free matrix factorization approach and should not be interpreted as a formal ancestry model or phylogenetic reconstruction. The components identified here represent co-occurring SNV signatures within rs2596542-T carrier haplotypes. Their interpretation as regulatory axes is supported by LD-reversal enrichment, genomic localization, and tissue-specific cis-eQTL annotation, but the components themselves are not discrete ancestral lineages.

Second, MICA protein-level allelic variation was not directly assessed. The present analysis focused on haplotypic SNV structure and transcript-level regulatory effects. Therefore, possible contributions of MICA coding variation, protein expression, shedding, or soluble MICA levels remain outside the direct scope of this study.

Third, this study does not directly test the rs2596542 disease-association flip-flop itself. Instead, it characterizes the upstream haplotypic and regulatory architecture that could plausibly mediate such reversals. The connection between c5-driven HLA-C downregulation and population-differential HCC susceptibility therefore remains inferential. Direct testing would require ancestry-diverse cohorts with matched genotype, expression, and disease-outcome data.

Finally, the framework was applied primarily to a single MHC locus. As a proof-of-concept extension, reanalysis of the COMT Val158Met flip-flop example reproduced the signed-LD structure reported by Lin et al. in matched 1000 Genomes populations and showed that Val carriers occupy distinct latent haplotype backgrounds across populations. However, broader application to additional loci will be needed to determine the generality of the approach. Formal simulation-based evaluation was not performed. Given the complexity of extended haplotype architecture, multiple embedded regulatory axes, population-dependent mixture proportions, and LD decay, simplified simulations may be less informative than systematic empirical application to additional GWAS loci with known cross-population effect-direction inconsistency.

Together, these findings support a composite-tagging model in which rs2596542-T captures separable MICA-regulatory and HLA-B/HLA-C-regulatory haplotypic backgrounds. This model provides a mechanistic explanation for cross-population signed-LD reversal at the locus and highlights the value of carrier-haplotype decomposition for interpreting complex MHC association signals.

## Funding

This research received no specific grant from any funding agency in the public, commercial, or not-for-profit sectors.

## Author contributions

Y.I. designed the study, performed all analyses, and wrote the manuscript.

## Declaration of interests

The authors declare no competing interests.

## Data and code availability

1000 Genomes Phase 3 VCFs from IGSR. GTEx v8 eQTL summaries from gtexportal.org. Analysis code: https://github.com/mountbook-lab/mica-rs2596542-haplotype-decomposition, archived at Zenodo (https://doi.org/10.5281/zenodo.20524043). Coordinates in GRCh37/hg19.

## Declaration of generative AI and AI-assisted technologies in the manuscript preparation process

During the preparation of this work, the author used OpenAI ChatGPT, OpenAI Codex, Anthropic Claude, and Claude Code to assist with language editing, manuscript organization, drafting support, and code review. After using these tools, the author reviewed and edited the content as needed and takes full responsibility for the content of the manuscript.

## Notes

https://zenodo.org/records/20524043

### Competing Interest Statement

The authors have declared no competing interest.

### Summary of Updates

This revision reorganizes the manuscript and supplementary materials, removes LIRI-related analyses reserved for a separate study, updates supplementary figure/table numbering and the Zenodo archive DOI, and corrects supplementary figure labels. The main conclusions are unchanged.

