## Supplementary Notes and Figures for "Carrier-set topology reveals composite MHC tagging at the MICA rs2596542 locus: insights from a flip-flop association example"

### Supplemental Notes

#### *Supplemental Note S1: Reference-panel replication and NMF extension of the COMT Val158Met flip-flop example*

As an external reference-panel replication and extension, we revisited the COMT Val158Met example from Lin et al. using 1000 Genomes Phase 3 phased haplotypes. rs4680 (chr22:19,951,271 G>A, Val158Met) and the P2 promoter marker rs2097603 (chr22:19,928,092 G>A) were both identified in the 1000 Genomes Phase 3 call set after coordinate-based lookup. Allele coding followed Lin et al.: A = Val at rs4680 (reference allele) and B = rs2097603 ALT. This coding specification is essential for sign reproducibility of signed LD values.

The signed correlation between rs4680 Val and rs2097603 ALT reproduced Lin et al.'s reported direction in matched reference populations: JPT for Japanese (Lin r = +0.31; 1000G r = +0.559), CEU for Irish-like European ancestry (Lin r = +0.48; 1000G r = +0.483), and YRI for Yoruba (Lin r = +0.03; 1000G r = +0.047). Quantitative agreement was closest for CEU and YRI, whereas the JPT estimate was stronger but directionally concordant. No direct Cambodian analogue is present in 1000 Genomes; Lin et al. reported r = −0.22 for Cambodians. These results confirm that the signed-LD structure underlying the Lin flip-flop example is directionally reproducible in independent reference data.

Notably, signed r varied even within AFR populations. MSL (r = −0.091) and GWD (r = −0.086) showed weak negative values, whereas YRI, ESN, LWK, ACB, and ASW remained positive (r = +0.04 to +0.13). This intra-super-population sign reversal indicates that Lin-like signed-LD reversal can occur at a finer population scale than inter-continental comparisons, consistent with the main rs2596542 analysis in which inclusion of AFR populations increased the fraction of partner SNVs showing signed-r reversal. Unsigned $r^2$ obscured these directional differences.

Local NMF at $k = 4$, selected by reconstruction-error saturation, decomposed rs4680 Val-carrier haplotypes into four components: comp2, which was AFR-specific; comp3, which was EAS-enriched; comp4, which was EUR/SAS-enriched; and comp1, which represented a flatter background. Val-carrier component loadings showed population-dependent latent background assignment: AFR Val carriers mapped predominantly to comp2, EAS and AMR Val carriers to comp3, and EUR Val carriers to a comp3 + comp1 mixture. Although comp4 was enriched in EUR/SAS haplotypes in the broader local decomposition, rs4680 Val-carrier haplotypes in EUR were assigned primarily to comp3 + comp1, with comp4 relatively depleted. Thus, the same physical allele tagged distinct latent haplotype backgrounds across populations, demonstrating component-aware interpretation of a canonical marker allele outside the MHC.

Compared with the MHC rs2596542 analysis, the COMT region yielded a simpler decomposition, with components corresponding more closely to population background than to functionally distinct regulatory axes. This contrast likely reflects differences in LD-block internal diversity. The MHC, shaped by balancing selection and recombination suppression, harbors sufficient within-carrier-set haplotypic complexity for NMF to resolve functionally distinct regulatory signatures, whereas the COMT region has a simpler haplotype landscape in which NMF components converge more closely on population-level structure. Together with the signed-LD replication of Lin et al., this NMF extension demonstrates that the carrier-set decomposition framework can be applied outside the MHC, although the depth of functional interpretation depends on the underlying haplotypic diversity.

Supplementary Figure S4 summarizes the COMT reference-panel extension, including population-conditional signed LD and asymmetry C for rs4680–rs2097603, rs4680 allele-conditional k = 4 NMF component mixtures, and population-level COMT NMF mixture across all haplotypes.

##### *Supplemental Note S2: Component label correspondence and cross-run reproducibility*

For cross-referencing with the companion study, the eight k = 8 NMF component indices used here (c0–c7) map to that study's unified functional labels as follows: c4 → MICA-reg (Axis I); c6 → MICA-reg (Axis I), softly determined across runs (see below); c5 → HLA-BC-variable (Axis II); c1 → HLA-B-stable (B*57:01); c3 → MICB-INS/EAS; c7 → backbone; and c0 and c2 are concordant across runs but functionally uncharacterized. The c-indices are run-specific identifiers, whereas the functional labels are the portable names shared between the two papers.

Beyond the 50-seed reproducibility reported in the main text (Supplementary Figure S3, Table S6), the eight-component basis also reproduced across an independent NMF run computed on a different variant window and filter (companion study; window chr6:31.25–31.55 Mb with 6,276 carrier-MAF ≥ 0.01 SNVs, versus the ±250 kb / 7,116-SNV window used here). Both runs derive from the same 1000 Genomes Phase 3 phased haplotypes and the same 2,111 rs2596542-T carriers, so allele orientation required no harmonization. H-matrix signatures were restricted to the 4,873 SNVs shared between the two filtered sets and matched by Hungarian assignment on cosine similarity (identical assignment under Pearson). All eight components found a one-to-one cross-run partner with no run-specific component (cosine 0.71–0.99). The MICB-INS/EAS component (c3) matched at cosine 0.96 and was independently anchored by rs115231984, the biallelic MICB-INS proxy, reaching maximum loading on the same component in both runs; the backbone component (c7) matched at 0.99 and the HLA-B-stable component (c1; anchored by rs2395029) at 0.75. The HLA-B/HLA-C-variable axis (c5) and the secondary MICA-reg component (c6, which the companion study associates with a European haplotype background) formed a partially entangled pair across runs (cosine 0.71–0.72); none of the c5 (Axis-II) top-5% signature SNVs fell within the shared window, so its reported cross-run cosine is a lower bound and Axis II is validated within each run individually rather than across runs.

Supplementary Figure S1. NMF vs LDA decomposition on the same rs2596542-T carrier haplotype matrix (k = 8).

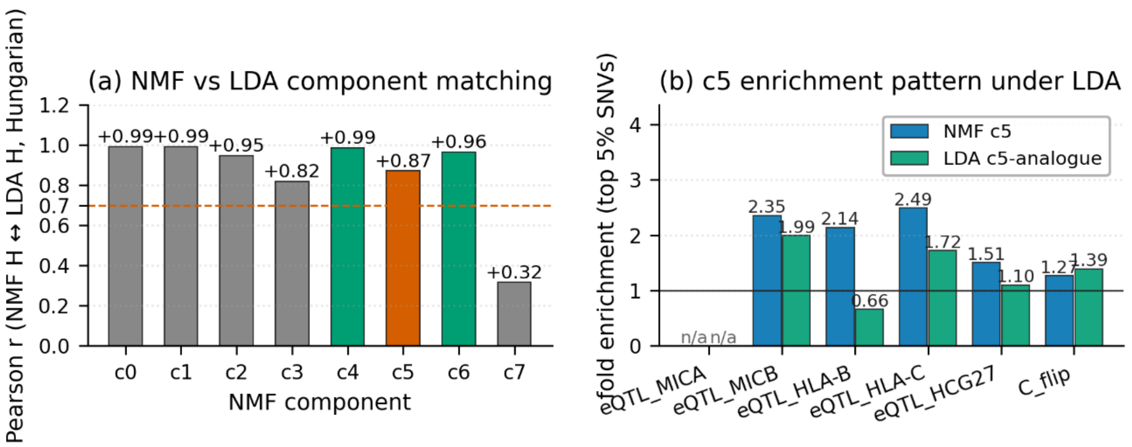


Supplementary Figure S2. NMF rank selection across k = 2–12.

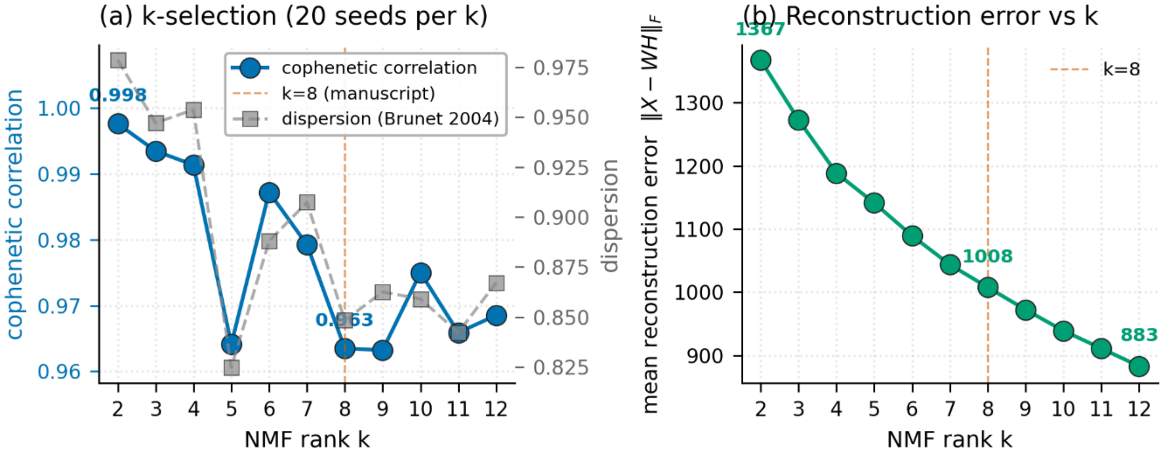

Supplementary Figure S3. NMF k = 8 reproducibility across 50 random initialisations.

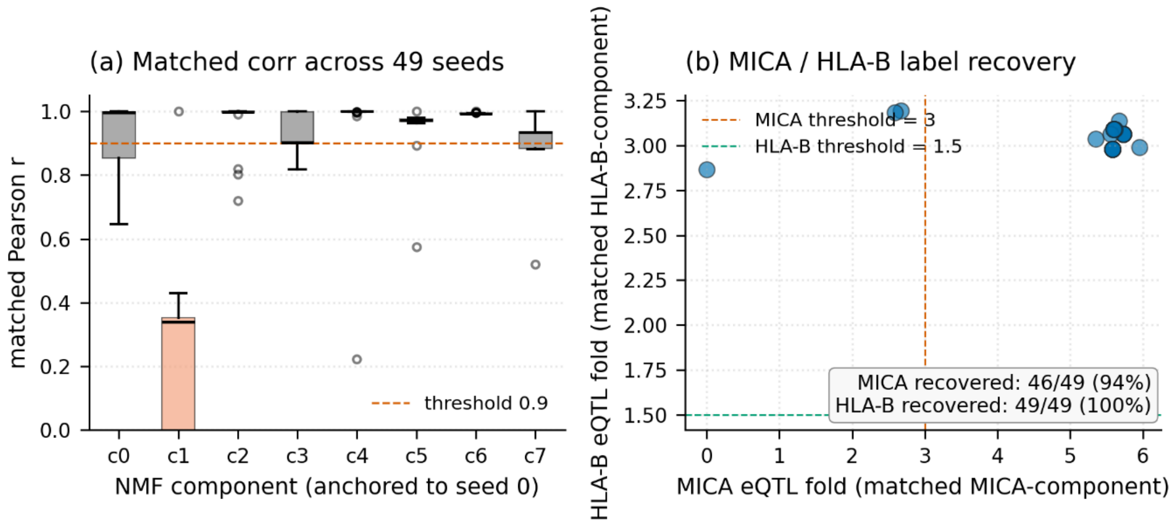


**Supplementary Figure S4. Reference-panel replication and NMF extension of the COMT Val158Met (rs4680) flip-flop example**

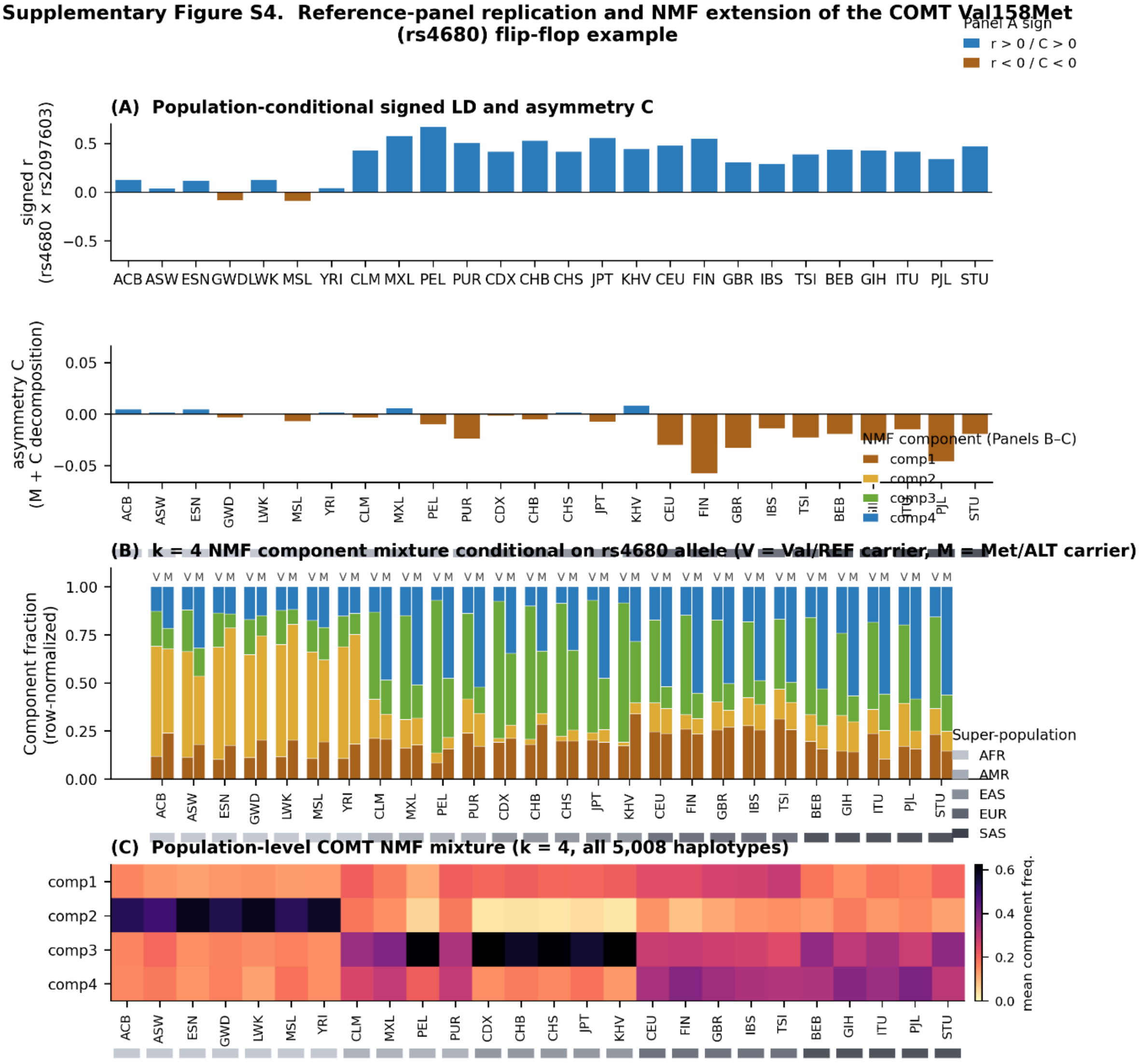

***Supplementary Table S1. Top-signature SNVs defining NMF components in rs2596542-T carrier haplotypes.*** *Top-signature SNVs were defined as variants with H-matrix loadings in the top 5% of each k = 8 NMF component. The table lists component-defining SNV sets used for LD-reversal enrichment, cis-eQTL annotation, matched-permutation analysis, and tag-SNP component mapping. The anchor SNV rs2596542 was removed from the NMF input matrix after carrier selection because all retained haplotypes carried the rs2596542-T allele.*

***Supplementary Table S2. MAF × distance matched permutation results.*** *Observed overlaps, matched-null expectations, fold enrichments, and z-scores for LD-reversal and gene-level eQTL test sets across NMF components.*

***Supplementary Table S3. GTEx tissue eQTL NES summary.*** *Per-tissue normalized effect-size summaries for component–gene eQTL overlaps across the six GTEx tissues used in the analysis.*

***Supplementary Table S4. LDA vs NMF comparison.*** *Comparison of c5-like enrichment profiles between NMF and LDA decompositions on the same rs2596542-T carrier haplotype matrix.*

***Supplementary Table S5. NMF rank selection.*** *Consensus cophenetic correlation, dispersion, and reconstruction-error summaries across NMF ranks used to select k = 8 as the primary rank.*

***Supplementary Table S6. NMF seed stability.*** *Hungarian-matched component correlations and label-recovery summaries across random initializations at k = 8.*

***Supplementary Table S7. Tag SNP component mapping.*** *Component-loading profiles and assignments for previously reported tag SNPs and proxies, including rs2244546 and rs2395029.*

***Supplementary Table S8. Axis sign coherence summary.*** *Direction counts, sign coherence, and mean NES summaries for the principal component–gene axes across GTEx tissues.*
